# p16-dependent upregulation of PD-L1 impairs immunosurveillance of senescent cells

**DOI:** 10.1101/2023.01.30.524522

**Authors:** Julia Majewska, Amit Agrawal, Avi Mayo, Lior Roitman, Rishita Chatterjee, Jarmila Kralova, Tomer Landsberger, Yonatan Katzenelenbogen, Tomer Meir-Salame, Efrat Hagai, Nemanja Stanojevic, Ido Amit, Uri Alon, Valery Krizhanovsky

**Author notes:** These authors contributed equally to this work (Julia Majewska, Amit Agrawal).

## Abstract

The accumulation of senescent cells promotes aging, but a molecular mechanism that senescent cells use to evade immune clearance and accumulate remains to be elucidated. Here, we report that p16-positive senescent cells upregulate the immune checkpoint protein programmed death-ligand 1 (PD-L1) to accumulate in aging and chronic inflammation. p16-mediated inhibition of CDK4/6 promotes PD-L1 stability in senescent cells via the downregulation of ubiquitin-dependent degradation. p16 expression in infiltrating macrophages induces an immunosuppressive environment that can contribute to an increased burden of senescent cells. Treatment with immunostimulatory anti-PD-L1 antibody enhances the cytotoxic T cell activity and leads to the elimination of p16, PD-L1-positive cells. Our study uncovers a molecular mechanism of p16-dependent regulation of PD-L1 protein stability in senescent cells and reveals the potential of PD-L1 as a target for treating senescence-mediated age-associated diseases.

## Main text

Senescent cells accumulate in tissues with age and disrupt tissue homeostasis, promote aging and limit lifespan (*1*). Conversely, the elimination of senescent cells significantly delays the onset of age-related phenotypes and extends lifespan in animal models (*2, 3*). However, the therapeutic potential of senolytic drugs that selectively clear senescent cells in humans is hampered due to their insufficient specificity and toxicity (*4*). Understanding the molecular mechanism that senescent cells use to impair their immunosurveillance, and thus allow their presence in aging tissues could provide new strategies for the regulation of senescent cell turnover.

Lines of evidence suggest that the age-dependent decline of immune-mediated clearance of senescent cells depends on senescent cell-autonomous mechanisms that inhibit their removal (*5, 6*), and functional impairment of the aged immune system (immunosenescence) (*7, 8*). Still, however, the intrinsic properties of senescent cells promoting their immunosurveillance escape and mechanisms explaining the immunosenescence-associated increased senescent cell burden in aging tissues or upon chronic damage remain to be elucidated.

## In aging p16-positive senescent cells upregulate immune checkpoint PD-L1

Immunosenescence has a causal role in driving systemic aging including an increased burden of senescent cells (*8*). To unravel cell-autonomous mechanisms leading to impaired immunosurveillance of senescent cells, we investigated functional phenotypes of immune cells undergoing senescence with age. We performed a mass cytometry analysis on lung immune cells from young 2-month- and aged 24-month-old C57BL/6 wild-type mice. viSNE analysis of the CD45^+^ immune cells distinguished interstitial and alveolar macrophages (IM and AM respectively), natural killer (NK) cells, eosinophils, neutrophils, CD4^+^ and cytotoxic CD8^+^ T cells, and CD11b^+^CD11c^+^ myeloid cells (Fig. 1A, fig. S1A). In aged lungs, the frequency of AM and CD11b^+^CD11c^+^ myeloid cells was significantly increased, with a distinct age-dependent infiltration of neutrophils (Fig. 1B, fig. S1B). This was accompanied by an age-associated significant reduction of NK cells, and a marked decrease of CD8^+^ T cells number in the line with the notion of age-dependent lung immunodeficiency (Fig. 1B) (*9*). We then analyzed identified immune cell types for the expression of senescence-related proteins. Remarkably, alveolar macrophages were particularly enriched for senescence (p16, p21), pro-inflammatory (p-p38, p-p65) and immunoregulatory (MHCI) markers in aged mice compared to young counterparts (Fig. 1C). Strikingly, AM also showed the highest expression of PD-L1 protein comparing to all other immune subsets (Fig. 1D). Cells expressing p16 are known to accumulate in aging tissues and promote age-related diseases (*3*). Interestingly, high-level expression of p16 in aged alveolar macrophages coincided with the significantly increased level of immune checkpoint molecule PD-L1 (Fig. 1E). Single cell analysis of AM also revealed a positive correlation between p16 and PD-L1 (Spearman’s rank correlation coefficient young - 0.9, old – 0.9; Fig. 1F). Consequently, we wanted to understand if p16 expression is coupled with the other proteins related to the senescence phenotype. We analyzed fold change of expression of senescence-related proteins as a function of p16 level and found that cyclin-dependent kinase inhibitor p21, anti-apoptotic protein BCL-XL and MHCI showed the most striking correlation with p16, irrespective of the age of mice (Fig. 1G). Increased expression of p21 and BCL-XL is known to underlie senescent cells’ resistance to apoptosis and can contribute to their accumulation in tissues (*10, 11*). To understand if the observed phenomena are not limited to AM, we analyzed lung epithelium, known to accumulate senescent cells in aging (*12*). Similarly, to AM, epithelial cells with a high level of p16 showed significant upregulation of PD-L1 expression in both, young and old mice (Spearman’s rank correlation coefficient young – 0.73, old - 0.86; Fig. 1H-I). We found out that expression of core proteins of senescence machinery (p21, p53), followed by proteins involved in inflammatory response (p-p65, p-p38) showed the strongest correlation with p16 level, regardless of age (Fig. 1J). These results indicate that the expression of p16 is correlated with PD-L1 expression in senescent cells, with higher levels of expression of both proteins in old mice. We hypothesized that macrophages with senescent phenotype might contribute to low-grade inflammation, but also induce a compensatory immunosuppressive response to prevent excessive tissue damage. Thus, in aging, increased expression of p16 associated with PD-L1-mediated immunosuppression, might lead to compromised immunosurveillance and consequently increased senescent cell burden.

**Fig 1.**
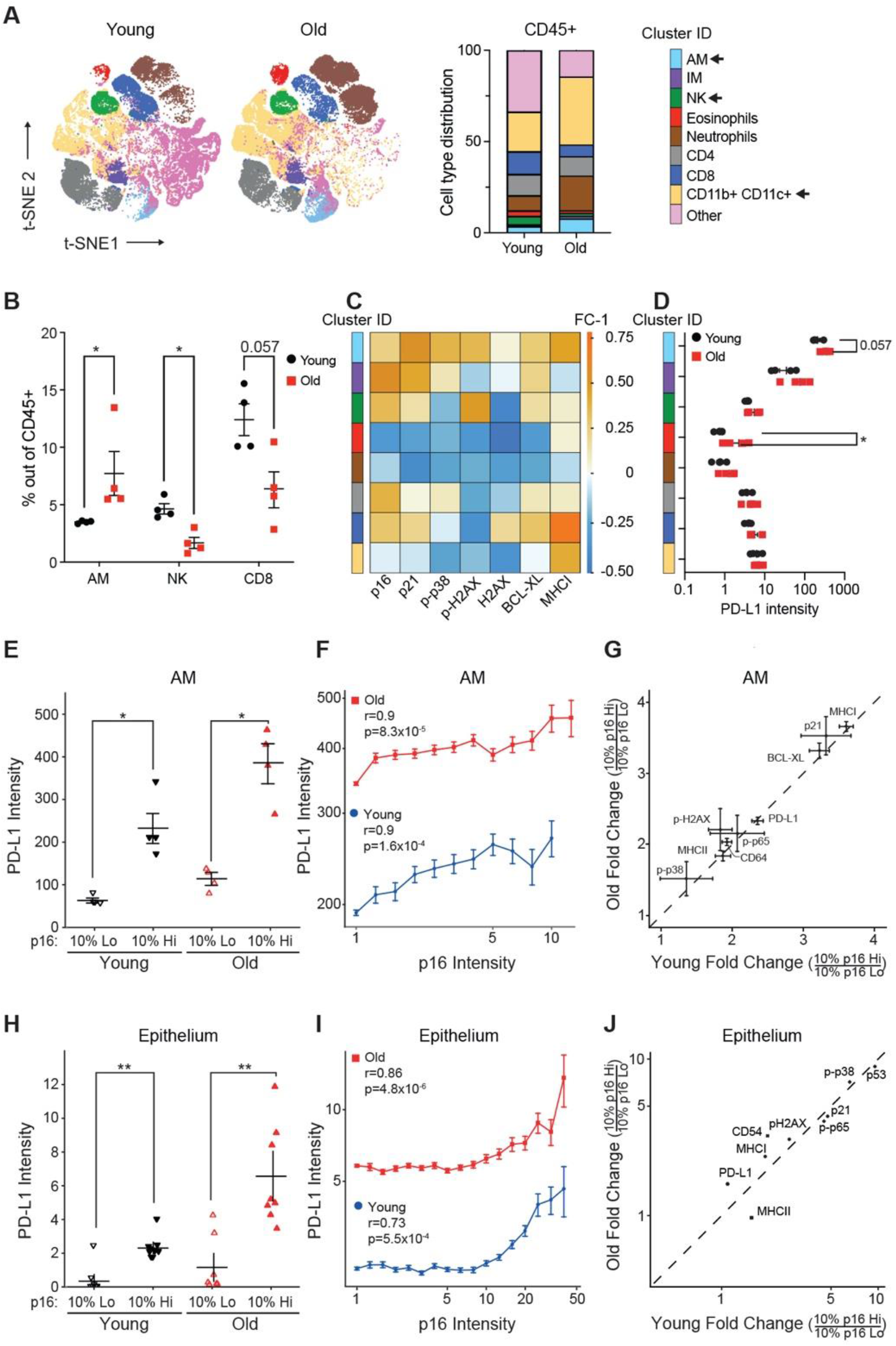
Expression of p16 is associated with PD-L1 upregulation in old mice. **(A)** Representative t-SNE plots of immune cell (CD45+) populations identified in 50,000 cells from all lung samples of old and young mice (left). Immune cell type distribution in young and old mice (right). Significantly changed populations are marked with an arrow (right). **(B)** The frequency of indicated immune cell types in young and old mice. **(C)** The heatmap of the change in the enrichment of senescence-related proteins in old mice for indicated immune cell types. **(D)** Comparison of mean expression of PD-L1 between identified immune cell populations. **(E)** Comparison of mean expression of PD-L1 between 10% highest and 10% lowest p16 expressing cells within alveolar macrophage population in young and old mice (n=4). **(F)** Spearman correlation between p16 and PD-L1 protein expression in alveolar macrophages in young and old mice. **(G)** Scatter plot showing fold change in expression of senescence-related markers between p16 high and p16 low expressing cells in alveolar macrophages in young and old mice. A diagonal line marks an equal fold change between young and old. **(H)** Comparison of mean expression of PD-L1 between 10% highest and 10% lowest p16 expressing cells within lung epithelium in young and old mice (n=8) **(I)** Spearman correlation between p16 and PD-L1 expression in lung epithelium in young and old mice. **(J)** Scatter plot showing fold change in expression of senescence-related markers between p16 high and p16 low expressing cells in the epithelium in young and old mice. **(F, I)** Spearman correlation coefficient (r) and associated p-value (p). Single cells were ranked by p16 expression level in bins from low to high. For each bin, the mean expression level of PD-L1 is shown. Alveolar macrophages (AM), Interstitial Macrophages (IM). Whitney-Mann U test was used for statistical analysis. *p < 0.05, **p < 0.01, ***p < 0.001

## In chronic inflammation p16-positive senescent cells upregulate immune checkpoint PD-L1

Accumulation of senescent cells promotes chronic inflammation, one of the most prominent age-related tissue phenotypes (*13*). We, therefore, asked if p16-positive senescent cells upregulate PD-L1 in chronic lung inflammation, which could explain their compromised immune clearance. To address this question, we used a mouse model of LPS-induced chronic lung inflammation, which resembles the pathology of chronic obstructive pulmonary disease (COPD) (*14*). Previously, we have reported that senescence of lung epithelium is implicated in COPD pathology and promotes chronic lung inflammation (*15*). Thus, to comprehensively characterize senescence in the epithelium, we took an unbiased mathematical approach to study cellular and phenotypic diversity of p16-positive cells within the non-immune, non-endothelial cell compartment (CD45^−^CD31^−^) in chronic lung inflammation by mass cytometry. Principal component analysis (PCA) of the data identified three distinct clusters (designated as clusters 1, 2, and 3) within epithelium derived from inflamed (LPS, Infl) and control (PBS) lungs (Fig. 2A). Density of cells within cluster 1, but not the other two clusters, significantly increased upon chronic inflammation (Fig. 2B). Cells within cluster 1 were significantly enriched in the expression of epithelial marker EpCam, and other canonical markers of lung epithelium subsets (SPC/SPB – alveolar cell type II, HOPX/PDPN – alveolar cell type I, CC10 – club cells; Muc5AC – goblet cells; KRT5/KRT14 – basal cells; KRT8 - alveolar progenitor cells, Fig. 2C, left, fig. S2B), indicating their epithelial identity. While cells in cluster 2 showed expression of basal progenitor cells (KRT14) (fig. S2B), cluster 3 was enriched in fibroblast markers (CD90.2, CD140a, CD140b) (Fig. 2C, right). Analysis of senescence markers of cells in cluster 1 showed enrichment for proteins of cell cycle arrest (p16, p21, p53), and depletion of the proliferation marker Ki-67 (Fig. 2D). Cells in cluster 1 were also enriched for molecules involved in the pro-inflammatory pathways (p-p38, p-p65, CD54), which can promote chronic inflammation (Fig. 2D). Remarkably, cells in cluster 1 also upregulated molecules associated with antigen presentation (MHCI, MHCII) and the immune checkpoint molecule PD-L1 (Fig. 2E), similar to what we observed in the lungs of old mice. These results suggest that p16-positive senescent cells express PD-L1, thus providing an intrinsic mechanism for their escape from immunosurveillance, and explaining their role in chronic inflammation by dysregulating immune responses.

**Fig 2.**
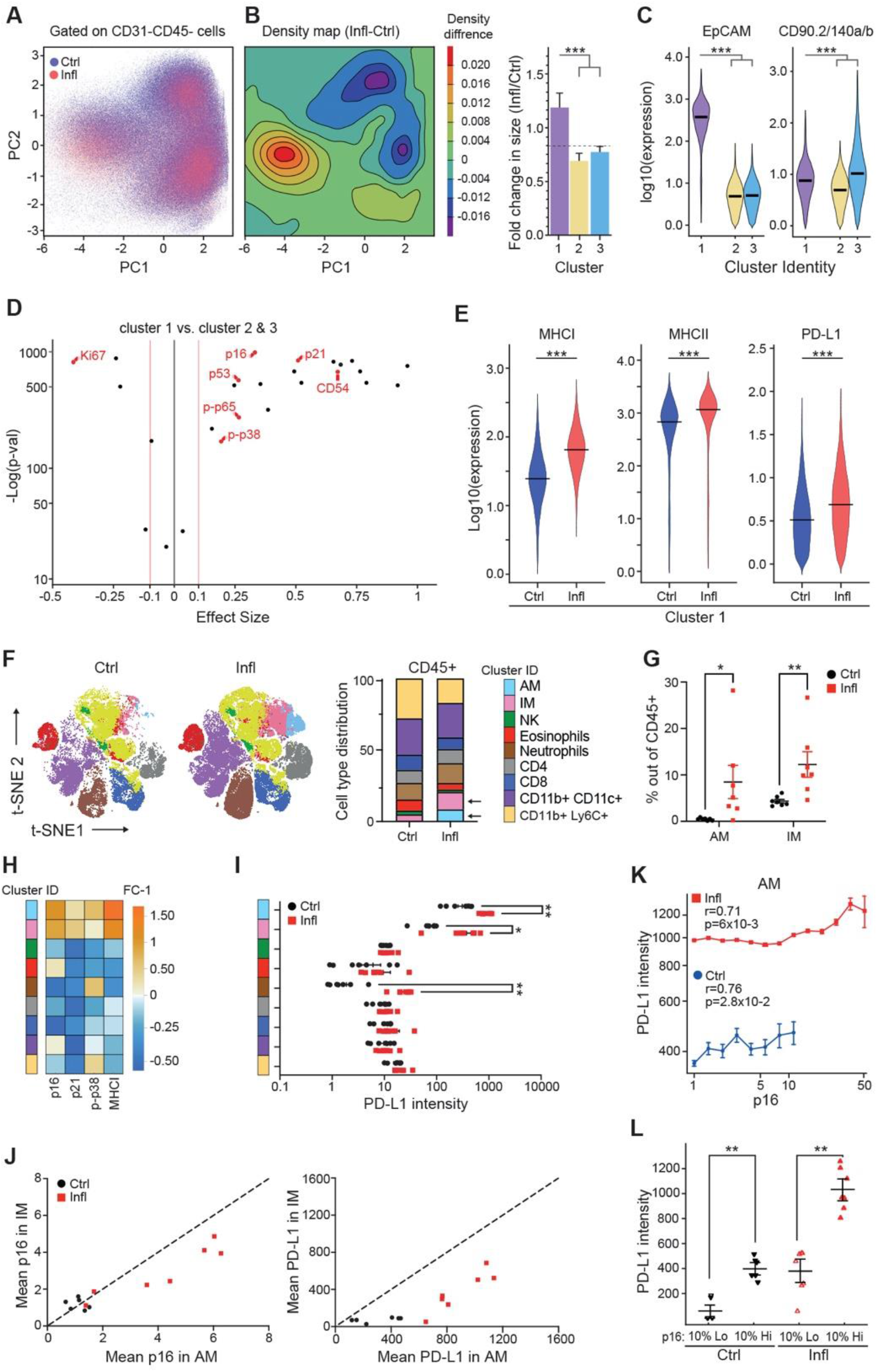
Expression of p16 is associated with PD-L1 upregulation during chronic inflammation. **(A)** Principal component analysis of CD45-CD31-cells from control (PBS, Ctrl) and chronically inflamed (LPS, Infl) lungs identifies three clusters (1, 2, and 3) based on their PC values (n=4-5). **(B)** Cell density map for each cluster shown as a difference in Kernel density distribution between Ctrl and Infl condition. Quantification of the fold change of cell frequency for each cluster (right). Error bars were estimated by bootstrapping. 10^4 cells were sampled from each cluster with sampling repeated 10^4 times. **(C)** Violin plot displaying the expression intensity distribution of epithelial marker (EpCAM) and fibroblast markers (CD90.2, CD140a, and CD140b) for identified clusters. Statistical significance was calculated by the Kruskal-Wallis test. **(D)** Volcano plot displaying enrichment of senescence-related proteins and depletion of the proliferation marker Ki-67 within cluster 1 in comparison to clusters 2 and 3. **(E)** Violin plot displaying the expression intensity distribution of the indicated proteins between Ctrl and Infl mice within cluster 1. **(F)** Representative t-SNE plots of immune cell (CD45+) populations identified in 50,000 cells from all lung samples of Ctrl and Infl mice (left). Immune cell types distribution in Ctrl and Infl mice. Significantly changed populations of alveolar (AM) and interstitial (IM) macrophages are marked with an arrow (right). **(G)** Frequency of AM and IM in the lungs of Ctrl and Infl mice. **(H)** The heatmap of the change in the enrichment of senescence-related proteins in Infl mice for indicated immune cell types. **(I)** Comparison of mean expression of PD-L1 between identified immune cell populations. **(J)** Comparison of mean p16 (left) and PD-L1 (right) expression between AM and IM. **(K)** Spearman correlation coefficient (r) between p16 and PD-L1 expression in AM of Ctrl and Infl mice and associated p-value (p). Single cells were ranked by p16 expression level in bins from low to high. For each bin, the mean expression level of PD-L1 is shown. **(L)** Mean PD-L1 expression in 10% highest and 10% lowest p16 expressing cells within AM in lungs of Ctrl and Infl mice (n=4). Mann-Whitney test was used for statistical analysis unless otherwise noted. *p < 0.05, **p < 0.01, ***p < 0.001

We aimed to understand how epithelial cells mediate the dysregulation of immune responses upon chronic damage. To address this, we performed RNA-sequencing of purified lung epithelial cells from LPS-treated and control mice. Gene ontology (GO) enrichment analysis of genes upregulated in the lung epithelium of mice with chronic inflammation indicated a marked increase in the expression of genes involved in a pro-inflammatory response, antigen processing, and presentation (fig. S2C-D). Furthermore, we observed an increase in gene signatures related to the activation of adaptive and innate immune responses, suggesting that the epithelial compartment enriched with senescence enhances the activation of macrophages during chronic inflammation.

We hypothesized that also chronic inflammation might promote immune cells to undergo senescence with compromised immune cell function causing a higher burden of damaged, or senescent cells. We evaluated immune cell populations using senescence and immune-cell centric antibody panel. viSNE analysis performed on CD45^+^ immune cells distinguished AM, IM, NK cells, eosinophils, neutrophils, CD4^+^ and CD8^+^ T cells, B cells, and CD11b^+^Ly6C^+^ myeloid cells (Fig. 2F; fig. S2E). In chronically inflamed lungs, we observed a significant increase in AM and IM (Fig. 2G). We analyzed identified immune cell populations for the expression of senescence-related proteins. Only AM and IM were enriched for the expression of senescence (p16, p21), pro-inflammatory (p-p38) and immunoregulatory (MHCI) markers in chronically inflamed compared to control mice (Fig. 2H; fig. S2F-G). The macrophages also showed the highest expression of PD-L1 protein compared to all other immune subsets, both in steady state and during inflammation (Fig. 2I). However, the mean expression levels of p16 and PD-L1 were higher in alveolar than in interstitial macrophages (Fig. 2J). Interestingly, high-level expression of p16 in AM co-related with significantly increased PD-L1 levels, consistent with our observation of age-related changes in AM (Spearman’s rank correlation coefficient LPS - 0.71, PBS – 0.76; Fig. 2K-L). Together, these results show that chronic inflammation triggered the accumulation of macrophages with a senescent phenotype and induced the expression of the immunosuppressive molecule PD-L1 in these cells.

Alveolar macrophages and the epithelial lining of the respiratory tract are the first lines of cellular and physical defense against environmental hazards such as cigarette smoke, allergens, and other air-born pollutants (*16, 17*). The pulmonary epithelium also interacts with immune cells to maintain homeostasis while facilitating immune response when necessary (*16*). Senescence of lung epithelium promotes chronic lung inflammation (*15*), and upregulation of immune checkpoint protein PD-L1 in both senescent epithelial cells and macrophages with senescent phenotype might be an intrinsic mechanism to evade immune clearance and promote persistent senescence in the tissue. We suggest that the upregulation of immune checkpoint PD-L1 on damage-induced senescent cells minimizes autoreactive immune responses in chronic inflammation to preserve organ integrity and function.

## p16-mediated inhibition of CDK4/6 stabilizes PD-L1 in cellular senescence

To explore the molecular mechanism behind the elevation of PD-L1 in senescent cells, we used a classical model of cellular senescence in primary human fibroblasts (IMR90) induced by DNA damage (*10*). Senescence resulted in a significant increase in surface PD-L1 expression compared to growing control cells (Fig. 3A). Replicative exhaustion, another physiological inducer of senescence also caused PD-L1 upregulation in these primary cells (Fig. 3A). Therefore, PD-L1 is elevated in senescent cells as an intrinsic response to stress, known to induce p16 expression, regardless of senescence-inducing stimulus or species.

**Fig 3.**
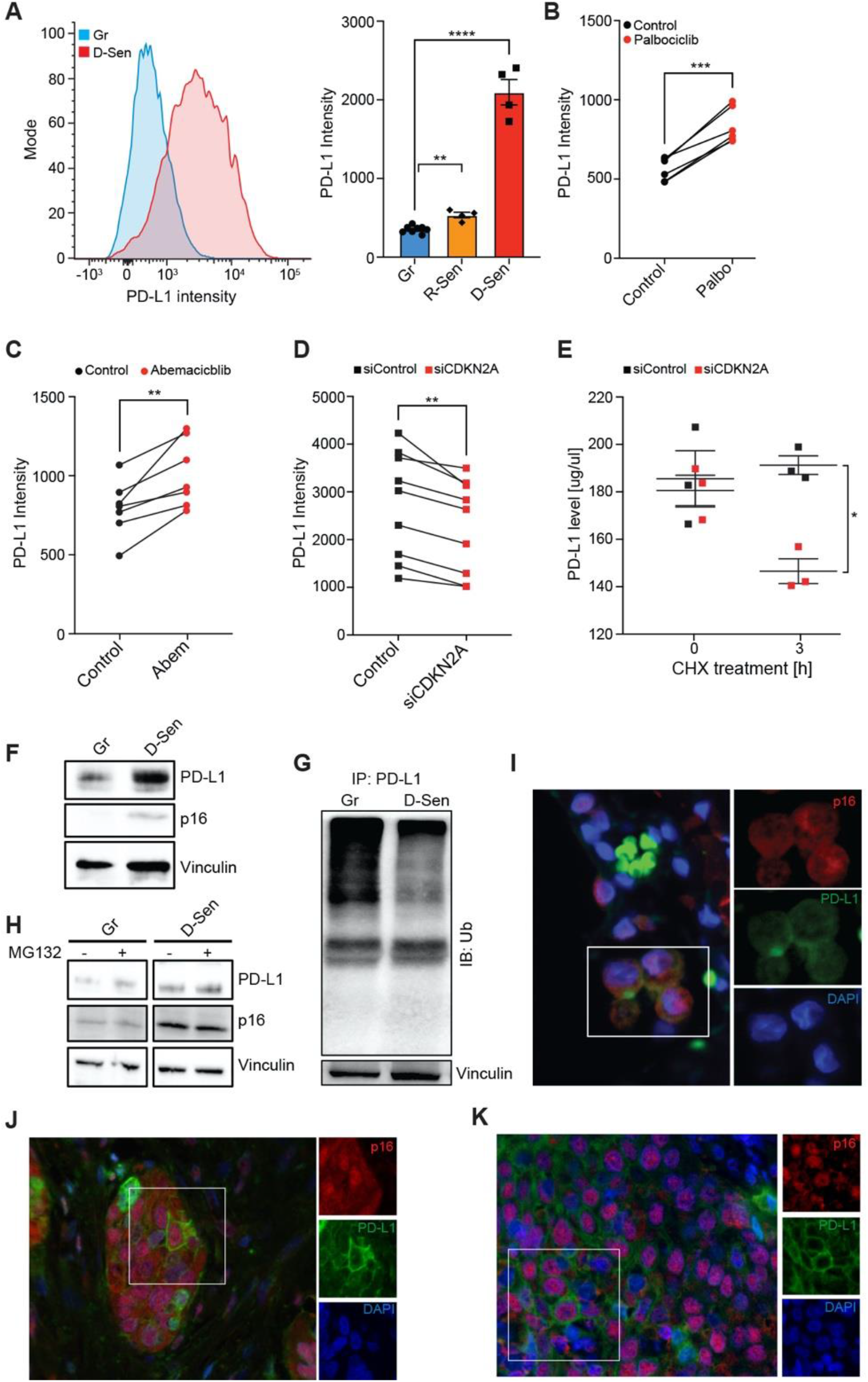
p16 regulates the stability of PD-L1 protein in senescent cells. **(A)** Flow cytometry analysis of PD-L1 expression in DNA damage-induced senescence (D-Sen) and Replicative senescence (R-Sen) in Primary Human Lung Fibroblasts (IMR-90) comparing to growing (Gr) IMR90 cells. Representative histograms (left) and quantification of median fluorescent intensity (MFI) of PD-L1 expression (right). **(B-C)** PD-L1 protein expression in growing IMR90 cells treated with 1 μM of CDK4/6 inhibitors (B) Palbociclib (Palbo) or (C) Abemaciclib (Abem) or equivalent amount of dimethyl sulfoxide (DMSO, control) for 48h. **(D)** PD-L1 protein expression in D-Sen treated with siControl or siCDKN2A. **(E)** ELISA-based measurement of PD-L1 protein levels in D-Sen treated with siControl or siCDKN2A and 200 μM cycloheximide (CHX) for 3h. **(F-H)** Immunoblot (IB) analysis of whole cell lysates derived from (F) Gr and D-Sen cells and (G) Gr and D-Sen cells treated with 10 μM MG132 or DMSO (negative control) for 3h. **(H)** Immunoblot analysis of Ubiquitin in immunoprecipitated PD-L1 protein derived from Gr and D-Sen cells. **(I-K)** Representative immunofluorescence images of p16 (red) and PD-L1 (green) staining in human pathologies: **(I)** emphysema, **(J)** lung adenocarcinoma, **(K)** squamous cell carcinoma patient. Blue, nuclei stain by DAPI. PD-L1 expression in A-D was quantified by flow cytometry analysis as median fluorescent intensity (MFI). (A) One-way ANOVA; n=3, Error bars, mean ± SEM. **p < 0.01, ****p < 0.0001, (B-E) Two-tailed unpaired Student t-test; n=3-9, Error bars, mean ± SEM. *p < 0.05, **p < 0.01, ***p < 0.001

p16 inhibits cyclin-dependent kinases (CDKs)4/6 to induce proliferative arrest. To test if the observed PD-L1 upregulation on senescent cells is triggered by p16-mediated inhibition of CDKs, we treated growing primary fibroblasts with highly selective inhibitors of CDK4 and CDK6 (CDK4/6i), abemaciclib or palbociclib, which can be considered functional p16 mimetics (*18*). Treatment with CDK4/6i significantly elevated surface PD-L1 expression (Fig. 3B-C). To understand whether the observed increase of PD-L1 protein in senescent cells is mediated by p16-dependent inhibition of CDK4/6, we silenced gene encoding p16 (CDKN2A) in senescent cells. We found that CDKN2A knockdown efficiently reduced PD-L1 protein level in senescent cells (Fig. 3D). Since inhibition of proteasomal-mediated degradation induces stability of many proteins to ensure cellular integrity in response to stress (*19*), we hypothesized that this mechanism might be responsible for p16-mediated PD-L1 upregulation in senescent cells. To examine the effect of p16 on the stability of endogenous PD-L1 protein in senescent IMR90 cells, we used a cycloheximide (CHX) chase assay to block translation and measure the degradation rate of PD-L1 protein. Knockdown of p16 caused a significant acceleration of PD-L1 turnover 3h post CHX treatment (Fig. 3E), indicating that PD-L1 degradation is p16-dependent. These results demonstrate that p16-mediated inhibition of CDK4/6 promotes PD-L1 protein stability in cellular senescence.

E3 ubiquitin ligases control PD-L1 protein levels through ubiquitin-dependent proteasomal-mediated protein degradation (*20*). SCF (SKP1-CUL1-F-box protein) E3 ubiquitin ligases constitute the largest family responsible for the turnover of key regulatory proteins of the cell cycle (*21*). β-TrCP is a substrate recognition component of SCF complex, also known to mediate PD-L1 ubiquitination for subsequent degradation (*20*). We reasoned that in cellular senescence with irreversible cell cycle arrest, the activity of SCF might be altered, thus affecting PD-L1 ubiquitination. We found that β-TrCP levels in senescent cells are downregulated compared to growing control cells (fig. S3A), possibly contributing to elevated PD-L1 level in senescence (Fig. 3F). Depletion of p16 in senescent cells restored β-TrCP level, possibly explaining the accelerated degradation of PD-L1 in senescent cells with p16 knockdown (Fig. 3D-E, fig. S3B). Given that SCF *ubiquitin* ligase via β-TrCP governs the ubiquitination and degradation of PD-L1, we evaluated its ubiquitination levels upon downregulation of β-TrCP in senescent cells. Immunoprecipitation of PD-L1 indicated a significantly lower ubiquitination signal in senescent cells compared to growing ones (Fig. 3G). Treatment of cells with the proteasome inhibitor MG132 significantly increased PD-L1 protein level in growing, but to a lesser extent in senescent cells, further supporting the role of the proteasome in the regulation of PD-L1 in senescent cells (Fig. 3H). These results indicate that p16 in senescence is linked to the upregulation of PD-L1 stability via decreased proteasome-mediated degradation.

Multiple mechanisms regulate PD-L1 at the transcriptional, posttranscriptional, and posttranslational levels (*22*). The pro-inflammatory secretome of senescent cells upregulates PD-L1 mRNA level through the JAK-STAT pathway (*23*). *In cancer cells*, cyclin *D-CDK4/6* dependent phosphorylation of SPOP/Cullin-3 ubiquitin ligase complex destabilizes PD-L1 *via* proteasome-mediated degradation, but inhibition of CDK4/6 significantly elevates PD-L1 level (*24*). In this line, the therapeutic potential of PD1-PD-L1 immune checkpoint therapy correlates with the p16 expression on tumor cells (*24, 25*). We have observed distinct clusters of cells expressing p16 and PD-L1 in both lung adenocarcinoma and squamous cell carcinoma (Fig. 3I-J). In patients with emphysema, a tissue manifestation of COPD, we identified p16, PD-L1 double-positive cells in alveolar space (Fig. 3K). These data suggest that p16-positive cells expressing PD-L1, which suppresses anti-tumor response, could also contribute to immune suppression in aging, including chronic lung disease. Together, our results link p16-mediated inhibition of CDK4/6 and ubiquitination pathways to the regulation of PD-L1 stability, which could lead to potential strategies to enhance the immune response against PD-L1 in senescent cells.

## p16-positive alveolar macrophages show immunosuppressive phenotype

We observed that cellular senescence triggers p16-mediated induction of PD-L1 in aging and chronic inflammation. So far, it is unknown if p16 expression contributes to compromised immune surveillance. Therefore, we decided to investigate the functional significance of p16-positive cells in this process, focusing on AM, as an example, due to their distinct accumulation in chronic lung inflammation (Fig. 4A). We used INs-Seq, a novel sequencing technology that combines intracellular labeling with transcriptomics to study p16 expressing cells (*26*). Gene set enrichment analysis revealed marked upregulation of central genes involved in extracellular matrix modeling or the response to retinoic acid in p16 high-expressing alveolar macrophages (Fig. 4B). Remarkably, p16 expression was negatively correlated not only with the DNA replication but also with immune responses, including dendritic cell activation or chemokine secretion (Fig. 4B). Alveolar macrophages contribute to respiratory tolerance via retinoic acid-mediated induction of Foxp3 expression in naïve T cells (*27*). In this line, in chronically inflamed lungs, we observed a significant increase of Foxp3+ Tregs enriched with marked PD1 expression (Fig. 4C-D). Interestingly, Tregs secrete regulatory cytokines, such as IL-4, and IL-10, which can modulate anti-inflammatory macrophage phenotype suggesting the cross-regulation of the immune response (*27*). Together, these results suggest that high-level expression of p16 in alveolar macrophages induces an immunosuppressive environment with compromised immune responses and most probably leads to non-resolving inflammation upon chronic damage.

**Fig 4.**
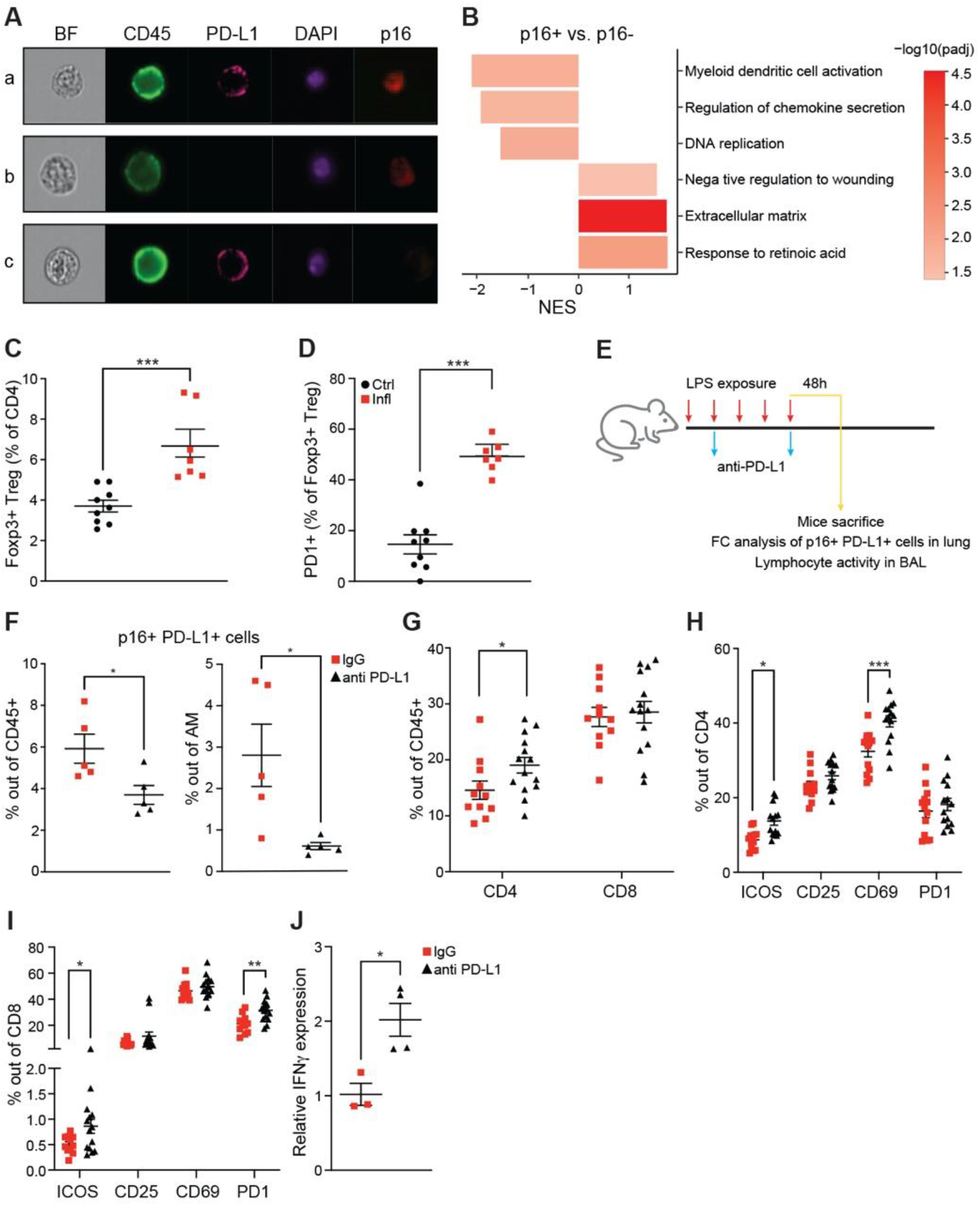
Anti-PD-L1 antibody treatment depletes p16, PD-L1-positive cells *in vivo*. **(A)** Imaging Flow Cytometry analysis of subcellular localization of p16 and PD-L1 staining within the alveolar macrophage (AM) population. Representative images of (a) CD45^+^PD-L1^+^p16^+^, (b) CD45^+^PD-L1^−^p16^+^, (c) CD45^+^PD-L1^+^p16^−^ cells. Bright-field (BF). **(B)** INs-Seq of p16^+^ AM. Gene Set Enrichment Analysis (GSEA) of p16^+^ and p16^−^ AM. **(C)** Quantification of Foxp3^+^ Tregs within CD4 population. **(D)** Quantification of CD4 Foxp3^+^ Tregs expressing PD1. **(E)** Experimental setup: mice exposed daily to either PBS (Ctrl) or LPS (Infl) inhalations for 5 days received an intravenous injection of anti-PD-L1 or matched IgG control as indicated, and analyzed 48h after the last inhalation. Mouse lungs and bronchoalveolar lavage (BAL) were harvested for subsequent analysis. **(F-I)** Flow cytometry analysis of lung **(F)** or BAL **(G-I)** from mice treated as in scheme E. **(F)** Percentage of p16^+^PD-L1^+^ cells within CD45^+^ or AM. **(G)** Percentage of CD4 and CD8^+^ T cells. **(H)** Percentage of CD4 T cells positive for ICOS, CD25, CD69, and PD1. **(I)** Percentage of CD8 T cells positive for ICOS, CD25, CD69, and PD1. **(J)** mRNA level of *IFN-gamma* cytokine from CD8^+^ T cells in the BAL. Values are relative to mice treated with IgG control. Quantification of indicated populations in C-D and G-I was performed by flow cytometry analysis. Bars correspond to median ± SEM; statistical significance was calculated using an unpaired student t-test. *p < 0.05, **p < 0.01, ***p < 0.001

Given the p16-dependent upregulation of PD-L1 in senescent cells, we wondered whether PD-L1 can serve as an extracellular marker to target senescent cells for depletion *in vivo*. To address this, we established a short-term LPS-mediated lung injury model, which results in infiltration of p16, PD-L1-double positive immune cells in LPS-stimulated mice compared to PBS control (fig. S4A-E). Specifically, alveolar macrophages identified as CD11c^+^SiglecF^+^ cells were enriched for PD-L1 and p16 expression, concomitantly with our results in the models of chronic inflammation and aging (fig. S4B; S4E). These experiments indicate that in response to stress, induction of endogenous p16 *in vivo* is associated with elevation of PD-L1 independently of the time scale of injury or identity of cell type responding to challenge. p16-mediated upregulation of PD-L1 in senescent cells makes it a potential target for monoclonal antibodies to stimulate anti-senescence immunity. Immunomodulatory PD-L1 antibodies can act either as antagonists to block the immune checkpoint PD1-PD-L1 axis or as agonists to enhance the immune response against target cells by engaging other immune system components (*28*). Engagement of activating Fcγ receptors on effector immune cells by anti-PD-L1 antibodies augments theirs *in vivo* activity (*29*) (fig. S4F). Therefore, we aimed to test if treatment with PD-L1-antibody activating Fcγ receptors on effector cells (activating PD-L1 antibody), could trigger an immune response and thus deplete p16^+^ cells from inflamed mouse lungs (Fig. 4E, fig. S4F). The treatment with activating anti-PD-L1 antibodies resulted in significant depletion of CD45^+^ positive for p16 and PD-L1, including a marked reduction of p16^+^PD-L1^+^ cells within the AM population, relative to the mice treated with isotype-matched control antibody (Fig. 4F). This experiment indicates that p16 expression renders cells permissive to depletion with anti-PD-L1 antibodies.

Stimulation of PD1 by PD-L1 limits effector functions and causes exhaustion in lymphocytes, mainly CD8 T cells. We tested if anti-PD-L1 antibody-mediated depletion of p16^+^PD-L1^+^ AM has a functional consequence for T cell activity. We decided to narrow the analysis of T cells to bronchioalveolar fluid, the functional lung compartment of AM. We tested activation markers on T cells isolated from inflamed lungs of mice treated with anti-PD-L1 antibody, or with isotype control. The activating anti-PD-L1 antibody treatment demonstrated significant efficacy in increasing the frequency of CD4 T cells showing co-expression of activation markers CD69 and ICOS (Fig. 4G-H). This response was accompanied by an anti-PD-L1-mediated increase in the percentage of CD8 T cells showing expression of activation markers ICOS and PD1, known to be expressed during early T cell activation (Fig. 4I). Activated CD8 T cells release IFN*γ* to enhance their cytotoxic function and motility (*30*). We have observed that anti-PD-L1 treatment also induced expression of *IFNγ* by CD8 T cells marking the activation of their effector function (Fig. 4J). Collectively, our data suggest that immunostimulatory PD-L1 antibody can enhance the cytotoxic effector function of T cells and elicit *in vivo* immune response against p16, PD-L1-double positive cells, leading to their depletion.

The axis between PD-L1 expressed on cancer cells and its receptor PD-1 on lymphocytes has a crucial role in suppressing anti-tumor immunity and serves as a major target in cancer immunotherapy (*31*). Both anti-PD-1 and -PD-L1 antibodies demonstrated therapeutic potential in multiple mouse models of cancer (*32, 33*) and human clinical trials (*34, 35*). The enhanced therapeutic function of PD-L1 antibodies can be achieved by the recruitment of effector cells with activating Fcγ receptors such as NK cells, macrophages, monocytes, and granulocytes (*36, 37*). We suggest that p16-positive senescent cells upregulate PD-L1 due to an increase in its stability and exploit the PD-1 pathway to promote immunosuppression and thus escape immune surveillance. It was recently shown that blocking PD-1 on T lymphocytes improves senescence surveillance (*38*). Here, we show accumulation of p16-positive cells with the senescent phenotype and elevated PD-L1 is associated with immunosuppression, possibly contributing to compromised immunosurveillance and increased senescent cell burden in aging tissues. Our data suggest that immunomodulatory PD-L1 antibody engaging Fcγ receptors on effector cells can be a promising approach to activate the immune response and deplete p16, PD-L1 double-positive senescent cells *in vivo*. Targeting PD-L1 in combination with other markers on senescent cells may offer new therapeutic opportunities to treat senescence-mediated age-associated diseases.

## Acknowledgments

We thank S. Jung for helpful advice and comments on the manuscript, N. Cohen for help with IV injections of anti-PD-L1 antibody, O. Regev for help with bronchoalveolar lavage, and S. Reich-Zeliger for stimulating discussions. V.K. is an incumbent of The Georg F. Duckwitz Professorial Chair.

## Funding

European Research Council grant 856487 (UA, VK)

Israel Science Foundation grant 2633/17; 1626/20 (VK)

Israel Ministry of Health 3-15100 (VK)

Weizmann – Belle S. and Irving E. Meller Center for the Biology and Aging (VK)

Irving E. Meller Center for the Biology and Aging (VK)

Sagol Institute for Longevity Research (VK)

## Author contributions

Conceptualization: JM, AA, VK

Methodology: JM, AA, AM, YK, TMS, EH

Investigation: JM, AA, AM, LR, RC, JK, TL

Visualization: JM, AM, TL

Funding acquisition: VK, UA, IA

Project administration: VK

Supervision: VK, UA, IA

Writing – original draft: JM

Writing – review & editing: JM, UA, VK

## Supplementary Materials

### Materials and Methods

#### Cell Culture

Human lung fibroblasts IMR-90 were purchased from ATCC (#CCL-186) and cultured to 70% confluency in Dulbecco’s modified Eagle (DMEM) medium supplemented with 10% fetal bovine serum (FBS) and 1% penicillin-streptomycin. To induce senescence, cells were treated with 50 μM etoposide (Sigma, #E1383) for 48h, washed 3 times with PBS, and cultured for additional 5-7 days in DMEM medium. Replicative senescence (RS) was induced by long-term passaging of the cells in tissue culture. Cells developed senescence phenotype after 35 population doublings. On the day of the experiment, cells were detached using trypsin.

#### siRNA

Cells were transfected overnight with 50nM of ON-TARGETplus SMARTpool siRNA targeting CDKN2A (#L-011007-00-0005) or with non-targeting siRNA pool (#D-001810-10-20) as control (Dharmacon). 24h post-transfection remaining adherent cells were harvested.

#### CDK4/6 inhibitors

Abemaciclib (Pubchem, #LY2835219) and palbociclib (Sigma, #PZ0383) were dissolved in DMSO (vehicle) to yield 10 mM stock solutions and stored at 80°C. IMR90 cells were treated with DMEM medium supplemented with either 1uM abemaciclib, palbociclib, or an equivalent amount of DMSO for 48h.

#### Proteasome inhibition

IMR90 cells were treated with DMEM medium supplemented with either 10 μM MG132 (Sigma, #M7449) or an equivalent amount of DMSO for 3h.

#### Immunoblot and immunoprecipitation assay

Cells were incubated in RIPA lysis buffer containing protease inhibitor cocktail (1:100) (Sigma, #P8340) and phosphatase inhibitor cocktail (1:100) (Sigma, #p5726) for 20 min on ice. Lysates were spun down for 15 min at 13,000 rpm and 4°C, and protein concentrations were determined with BCA assay (Thermo Scientific). Equal amounts of protein were resolved by SDS–PAGE and immunoblotted using β-TrCP (Cell Signalling Technology, CST-4394S), p16 (Abcam # Ab108349), vinculin (Abcam #Ab129002), PD-L1 (Cell Signalling Technology, #CST-13684), appropriate HRP-conjugated secondary antibody and ECL visualization.

For immunoprecipitations analysis, cells were lysed in HNTG buffer (0.05M Hepes pH 7.5, 10% Glycerol, 0.15M NaCl, 1% Triton X-100, 0.001M EDTA, 0.001M EGTA, 0.01M NaF, 0.025M β-glycerol phosphate) supplemented with protease inhibitor cocktail (1:100) (Sigma, #P8340) and phosphatase inhibitor cocktail (1:100) (Sigma, #P5726). 2,000 μg of total cell lysates were incubated with previously coated protein A/G agarose beads (Santa Cruz Biotechnology, #2003) with anti-PD-L1 antibody (3ug/ml) (Cell Signalling Technology, #13684) overnight at 4°C with gentle rotation. The beads were thoroughly washed with HNTG buffer and eluted with 6×SDS loading buffer by boiling at 95°C for 10 min. Ubiquitination of PD-L1 was measured by immunoblotting with anti-Ubiquitin antibody (Santa Cruz Biotechnology, #8017).

#### Cycloheximide Chase Assay

24h following transfection with siRNA, senescent IMR90 cells were treated with 200 μM cycloheximide (Sigma, #C4859) for 3h. Cells were lysed by incubation with 100 μl of RIPA buffer (supplemented with PMSF and protease inhibitor cocktail) for 20 min on ice and protein concentrations were determined using BCA assay. 10 μg of protein lysate was used to measure levels of PD-L1 by the enzyme-linked immunosorbent assay (ELISA), using the PD-L1/B7-H1 Quantikine ELISA Immunoassay kit (R&D, #DB7H10), according to the manufacturer’s protocol. The optical density of each well was measured with the Infinite 200 plate reader (Tecan) at 450 nm with wavelength correction set at 540 nm. The experiment was performed twice and each sample was performed in duplicate.

#### Immunofluorescent staining of human tissue microarray

FFPE sections of human lung tissue microarray (US Biomax, #LC487) were incubated at 60°C for 60 min, deparaffinized, and incubated in acetone for 7 min at −20°C, followed by subsequent incubation with 3% H_2_O_2_ for 15 min at room temperature to block endogenous peroxidase activity. Antigen retrieval was performed in a microwave (3 min at full power, 1000 W, then 20 min at 20% of full power) in Tris-EDTA buffer (pH 9.0). The slide was blocked with 20% NHS + 0.5% Triton in PBS and primary antibodies were diluted in 2% NHS + 0.5% Triton in PBS (PBST) (p16 1:30, Abcam #Ab108349; PD-L1 1:100, Abcam #Ab213524) in a multiplexed manner with the OPAL reagents (Akoya Bioscience), each one O.N. at 4°C. Following overnight incubation with the first primary antibody, the slide was washed with PBS, incubated in 2% NHS in PBS with secondary antibody conjugated to HRP (1:100) for 90 min, washed again, and incubated with OPAL reagents for 15 min. The slide was then washed and microwaved (as described above), washed, stained with the next primary antibody and with DAPI at the end of the cycle, and mounted. We used the following staining sequence: p16 → PD-L1→ DAPI. Each antibody was validated separately and then multiplexed immunofluorescence (MxIF) was optimized to confirm that the antibody signal was not lost or changed due to the multistep protocol. Slides were imaged with an Eclipse Ni-U microscope (Nikon), connected to a color camera (DS-Ri1, Nikon), and DAPI, Cy3, and Cy5 cubes. Images were analyzed using the Fiji software.

#### Mice

Mice 10 - 14 weeks of age (young) or 24 months old (old) were used in all experiments. Female C57BL/6 mice were purchased from Harlan Laboratories. All mice were housed and maintained under specific pathogen-free conditions at the Weizmann Institute of Science in accordance with national animal care guidelines. All animal experiments were approved by the Weizmann Institute Animal Care and Use Committee.

#### LPS exposure and Treatment

For chronic LPS exposure, mice were exposed to an aerosolized PBS alone or PBS containing Escherichia coli LPS (0.5 mg/ml; Sigma, #L2630) for 30 min, 3 times a week for 10 weeks, in a custom-built cylindrical chamber as described previously (*1, 2*). For short-term 5-day LPS exposure, mice were exposed as in chronic exposure, but only for 5 constitutive days. Mice were sacrificed and lungs were harvested 48h after the last exposure.

For PD-L1 antibody treatment, mice received an intravenous injection of 200 μg anti-PD-L1 (Ichorbio, #ICH1086), or 200 μg isotype control IgG2b (Ichorbio, #ICH2243) on the second and fifth day of short-term LPS inhalation.

BAL fluid was collected from perfused lungs by double washing with 1 mL PBS through a tracheal catheter as previously described (*1*).

#### Tissue Dissociation

To achieve single cell suspension from the lung, mice were euthanized by administration of xylazine/ketamine and then perfused by injecting cold PBS via the right ventricle before lung dissection. Lung tissues were dissected from mice, cut into small fragments, and suspended in 1.5 ml of Dulbecco’s modified Eagle medium/F12 medium (Invitrogen, #11330-032) containing elastase (3 U/ml, Worthington, #LS002279), collagenase type IV (1 mg/ml, Thermo Scientific, #17104019) and DNase I (0.5 mg/ml, Roche, #10104159001) and incubated at 37 °C for 20 min with frequent agitation. After the dissociation procedure, cells were washed with an equal volume of DMEM/F12 supplemented with 10% FBS and 1% penicillin–streptomycin (Thermo Scientific), filtered through a 100-μm cell strainer, and centrifuged at 380*g* for 5 min at 4 °C. Pelleted cells were resuspended in red blood cell ACK lysis buffer (Gibco, #A1049201), incubated for 2 min at 25°C, centrifuged at 380g for 5 min at 4°C and then resuspended in ice-cold sorting buffer (PBS supplemented with 2mM ethylenediaminetetraacetic acid, pH 8 and 0.5% BSA).

Flow cytometry. IMR-90 cells were stained with Zombie Aqua Viability fixable stain (#423101) for evaluation of live/dead cells, followed by antibody Brilliant Violet 711-PD-L1 (#329721) or isotype control (#400353) staining (all from Biolegend).

Lung single cell suspension was stained with anti-mouse CD16/32 (eBioscience, #14-0161-82) to block Fc receptors before labeling with fluorescent antibodies against cell-surface epitopes. For samples that were used for p16 intracellular staining, we used the following antibodies for extracellular staining: Brilliant Violet 605-CD45 (#103140), FITC-CD11c (#117306), Brilliant Violet 421-SiglecF (#155509) purchased from BioLegend. We used two clones of PD-L1 antibody (Brilliant Violet 785-PD-L1, 10F.9G2, #124331, PE-PD-L1, MIH6, #153611) purchased from Biolegend, which yielded similar results. Then cells were fixed with 90% methanol for 10 min at 4°C. All centrifugation steps after fixation were done at 850*g* for 5 min at 4 °C. For intracellular staining, cells were stained with p16 antibody (Abcam, #Ab54210) conjugated to Alexa Fluor 647 fluorophore (Thermo Scientific, #A20186). Cells were stained with Zombie Aqua Viability fixable stain for evaluation of live/dead cells. For the characterization of immune subsets in BAL we used the following antibodies: Pacific Blue-CD69 (#104523), Brilliant Violet 605-ICOS (#313537), Brilliant Violet 785-NK1.1 (#108749), PerCP-CD19 (#115531), FITC-CD3 (#100204), PE-CD25 (#102007), PE-Dazzle 595-TIGIT (#142109), PE-Cy5-CD8 (#100709), PE-Cy7-CTLA4 (#106313), APC-LAG3 (#125209), Spark Nir 685-CD4 (#100475), Alexa Fluor700-CD44 (#103025), APC/Cy7-PD1 (#135223), APC Fire810-CD45 (#103173) all from BioLegend. Cell populations were recorded using LSR-II new (BD Biosciences) or Aurora (Cytec), and analyzed using BD FACS DIVA software (BD Biosciences) and FlowJo software.

For imaging flow-cytometry cells were stained with FITC-CD45 (Biolegend, #103107), Brilliant Violet 786-PD-L1 (Biolegend, #124331), and Ax647-p16 (Abcam, #Ab54210, conjugated to Alexa Fluor 647 fluorophore from Thermo Scientific, #A20186). Before the acquisition, cells were stained with DAPI and filtered through a 100 μm membrane. Cells were acquired using ImageStreamX mark II (Amnis, Part of EMD Millipore Merck) and analysis of the image data was performed using IDEAS 6.2 software as described previously (*3*).

#### Mass Cytometry

All antibodies used in the study, their corresponding clone, provider, and catalog number are listed in Table 1. Antibodies were obtained in a protein-free buffer. Custom metal-conjugated antibodies were generated using MaxPAR antibody labeling kits (Fluidigm) or the MIBItag Conjugation Kit (IONpath) according to the manufacturer’s instructions. After metal conjugation, the concentration of each antibody was determined with a Nanodrop (Thermo Scientific), and adjusted to 0.5 mg/ml with Antibody Stabilizer PBS (CANDOR Bioscience, #131050) for long-term storage at 4°C.

Lung single cell suspension was washed once in 1 ml of Cell Staining Buffer (CSB) (Fluidigm, #201068). To ensure homogeneous staining, 4 x 10^6^ cells from each sample were used. For viability staining, cells were incubated with 1.25 μM Cell-ID Cisplatin (Fluidigm, #201064) for 3 min before quenching with CSB. Before antibody staining, cells were incubated for 10 min at 4°C with anti-mouse CD16/32 (Invitrogen, #14-0161-82) to block Fc receptors. Cells were stained with the epithelial or immune-centric antibodies for 45 min at 4°C. An antibody cocktail of extracellular markers was prepared as a master mix and 50 μl of the cocktail was added to the samples resuspended in 50 μl of CSB. Cells were washed twice with CSB and permeabilized with fixation/permeabilization buffer (eBioscience, #88-8824-00). Then, samples were washed twice with CSB, incubated with 5% goat serum (Sigma, #G-9023), and resuspended in 50 μl of CSB before the addition of 50 μl of a cocktail of intracellular antibodies. For DNA-based detection, cells were stained with 125 nM Cell-ID Intercalator-Ir (Fluidigm, #201192A) in PBS with 1.6% PFA (Electron Microscopy Sciences, #15700) overnight at 4°C. Cells were then washed once in CSB, and twice in Maxpar Water (Fluidigm, #201069). For mass cytometry acquisition, samples were diluted to 3 x 10^5^ cells/ml in Maxpar Water containing 1:10 EQ Four Element Calibration Beads (Fluidigm, #201078) and filtered through a 35-μm filter mesh tube (Falcon). For acquisition, the CyTOF Helios system (Fluidigm) was used and samples were acquired at the rate of 200 events/sec. Data was collected as .fcs files. Data were normalized, and concatenated when necessary, via the CyTOF software V7.0 (Fluidigm). Then, the Cytobank platform (Beckman Coulter) was used to gate out the normalization beads according to the 140Ce channel. Next, several gates were applied to gate out live cells for further analysis. First, live single cells were gated using the cisplatin 195Pt, iridium DNA label in 193Ir, followed by the event length, and the Gaussian parameters of width, center, offset, and residual channels.

To normalize data a hyperbolic arcsine transformation (with a scale factor of 5) was first applied. viSNE analysis and illustrations were generated using Cytobank software.

#### Flow cytometry cell sorting

Cell populations were sorted using BD FACSAria Fusion flow cytometer (BD Biosciences). Before sorting, all samples were filtered through a 70-mm nylon mesh. Populations that were sorted were epithelial cells (EpCam+ CD31-CD45-Ter119-), alveolar macrophages (CD45+CD11c+SiglecF+ p16 high/low), and CD8a T cells (CD45+SiglecF-CD3+CD8a+). Sytox Blue (Invitrogen, #34857) or Aqua Zombie was used for viability staining. 5,000-1000 live cells were sorted into a low-bind eppendorf tube containing 50 ul of lysis/binding buffer (Invitrogen). Immediately after sorting, samples were spun down, snap-frozen, and stored at −80°C until further processing.

To sort out lung epithelium, cells were stained with the following antibodies: Brilliant Violet BV605-CD31 (#102427), PE-CD45 (#103106), Alexa Fluor 488-EpCam (#118210) all purchased from BioLegend and eFluor450-TER-119 (eBioscience, #48-5921-82). To sort out alveolar macrophages, cells were stained with Brilliant Violet 605-CD45 (#103140), FITC-CD11c (#117306), Brilliant Violet 421-SiglecF (#155509), Brilliant Violet 786-PD-L1 (#124331) all from Biolegend and p16 (Abcam, #Ab54210) conjugated to Alexa Fluor 647 fluorophore (Thermo Scientific, #A20186).

To sort out CD8a T cells, cells were stained with Brilliant Violet 605-CD45 (#103140), Brilliant Violet 421-SiglecF (#155509), FITC-CD3 (#100204), APC-CD8a (#100711) all from BioLegend.

#### Preparation of libraries for RNA-seq

10^4^ cells of lung epithelium (EpCam+ CD31-CD45-) were sorted into 50 μL of lysis/binding buffer (Life Technologies). mRNA was captured with 15 μl of Dynabeads oligo(dT) (Life Technologies), washed, and eluted at 70°C with 6.1 μl of 10 mM Tris-Cl (pH 7.5). cDNA libraries were prepared from pooled samples of the same cell type (10000 cells per sample) according to a bulk variation of MARSseq (*4*) and were sequenced on Illumina NextSeq500 (Illumina). INs-seq libraries were prepared as previously described (*5*) followed by bulk MARS-sequencing.

#### RNA-seq analysis

Raw data were processed with the User-friendly Transcriptome Analysis Pipeline (UTAP) (*6*). Only reads with unique mapping to the 3’ of RefSeq annotated genes (mm10, NCBI Mus musculus Annotation Release 109) were analyzed. For differential gene expression analysis, we used DESeq2 (*7*), following standard workflow, to analyze RNA-seq count data derived from lung epithelial cells, comparing LPS to PBS. Genes with < 30 UMIs across samples were pre-filtered. Differentially expressed genes (DEGs) were selected to have |*fold* − *change*| > 1.25 and Benjamini-Hochberg adjusted *P* < 0.05. For Gene set enrichment analysis, we used DESeq2 (*7*) to derive gene fold-changes for LPS vs. PBS epithelial cells, and for p16+ vs. p16-macrophages, controlling for treatment (LPS/PBS) as a covariant. We then applied gene set enrichment analysis (GSEA) to the ranked fold changes. We used Fast Gene Set Enrichment Analysis (“fgsea”) library (*8*) implemented in R to test for enrichment of gene sets (#genes > 10) from the mouse C5 v5p2 gene ontology (GO) collection of the Molecular Signature Database (*9*).

#### Real-time RT–PCR analysis

mRNA was extracted from 5000 CD8a T cells sorted into 50ul of lysis/binding buffer (Invitrogen) and captured using Dynabeads oligo(dT) (Invitrogen) kit according to the manufacturer’s protocols. For qPCR analysis, mRNA was reverse transcribed using SuperScript II (Invitrogen, Cat#11904018), and cDNA was diluted 1:10 for qPCR analysis performed using the SYBR Green system. The relative gene expression was determined using the ΔΔCt method and normalization to *Actb*. We used four biological replicates for each condition. One-tailed t-tests were used to perform statistical analysis.

The following primers were used: mouse *Actb*: forward, 5’-GGAGGGGGTTGAGGTGTT-3’, reverse, 5’-TGTGCACTTTTATTGGTCTCAAG-3’; mouse *Ifng*: forward, 5’-TGAACGCTACACACTGCATCTTGG-3’, reverse, 5’-CGACTCCTTTTCCGCTTCCTGAG-3’.

#### Statistical analysis

Data are presented as means ±SEM unless otherwise noted. Comparisons between two groups were performed by an unpaired two-tailed Student’s t-test unless otherwise noted. Chi-squared test was performed for RNA-seq analysis of DEGs. For consistency in comparisons, significance in all figures is denoted as follows: \**P* < 0.05, \*\**P* < 0.01, \*\*\**P* < 0.001, \*\*\*\**P* < 0.0001.

**Fig. S1.**
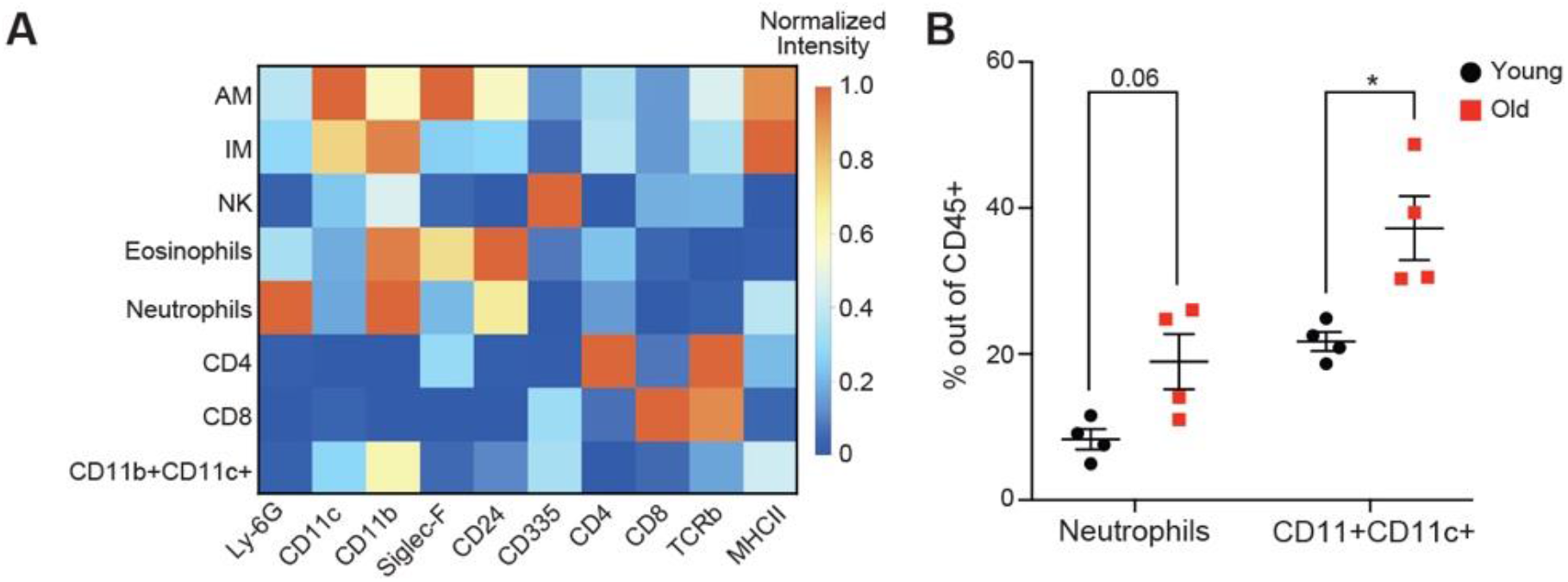
Expression of p16 is associated with PD-L1 upregulation in old mice. **(A)** The heatmap of normalized lineage markers expression for identified immune cell types. **(B)** Frequency of indicated immune cell types in young and old mice.

**Fig. S2.**
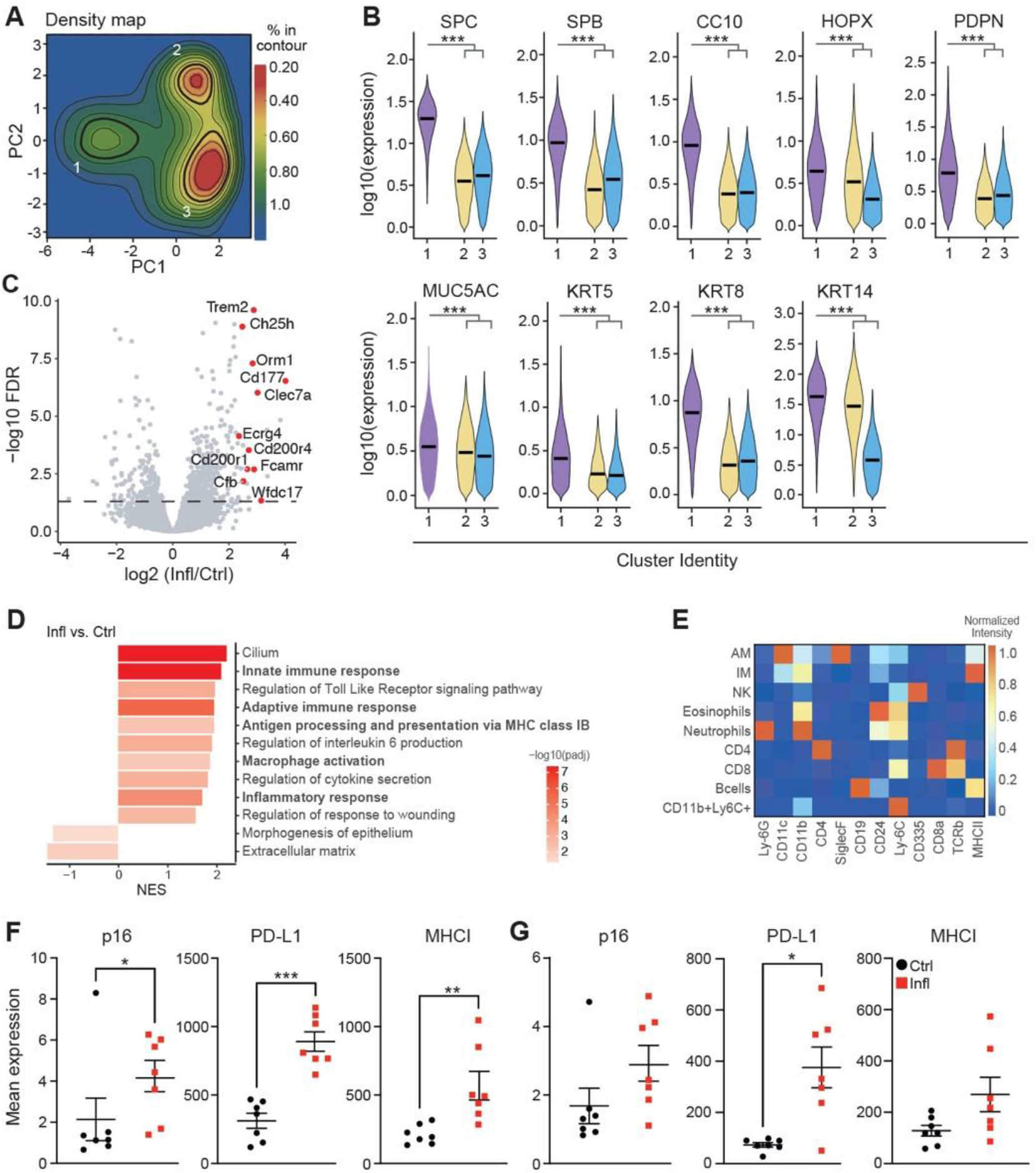
Expression of p16 is associated with PD-L1 upregulation during chronic inflammation. **(A)** Cell density map for each cluster as a function of Kernel density distribution in Ctrl and Infl condition. **(B)** Violin plot displaying the expression intensity distribution of phenotypic markers for identified clusters. Statistical significance was calculated by the Kruskal-Wallis test. **(C-D)** RNA-seq of lung epithelial cells derived from lungs of Ctrl and Infl mice **(C)** Volcano plot of differentially expressed genes (DEGs) between Ctrl and Infl mice (n=5). DEGs were identified by Padj < 0.05 and |Fold change| > 1.25. Immune response-related genes are labeled in red. **(D)** Gene Ontology (GO) enrichment analysis of DEGs identified in (C). GO terms are ordered by Normalized Enrichment Score (NES). A positive NES value indicates enrichment in Infl condition, negative NES value indicates enrichment in the Ctrl phenotype. The color of the bar denotes −log10 (p-value). **(E)** Heatmap of normalized lineage marker expression for indicated immune cell types. **(F-G)** Comparison of mean expression of p16, PD-L1, and MHCI markers within **(F)** AM and **(G)** IM in Ctrl and Infl mice (n=7). Mann-Whitney test was used for statistical analysis unless otherwise noted. *p < 0.05, **p < 0.01, ***p < 0.001

**Fig. S3.**
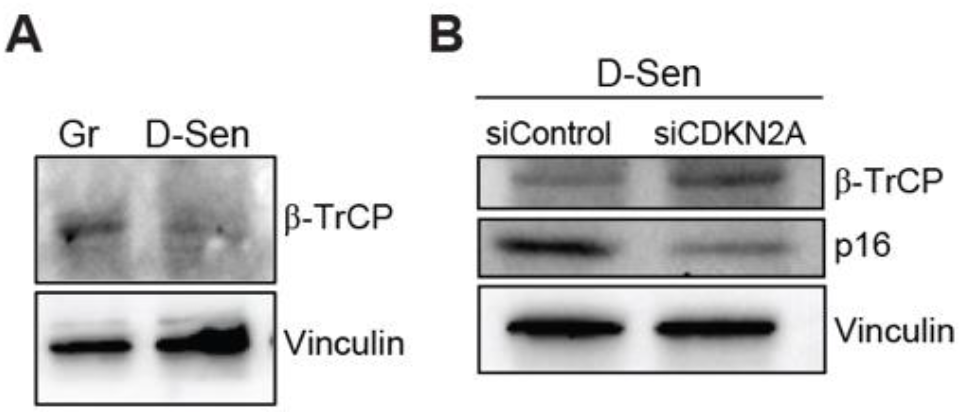
p16 regulates the stability of PD-L1 protein in senescent cells. **(A-B)** Immunoblot (IB) analysis of whole cell lysates derived from (A) Gr and D-Sen cells and (B) D-Sen cells treated with siControl and siCDKN2A

**Fig S4.**
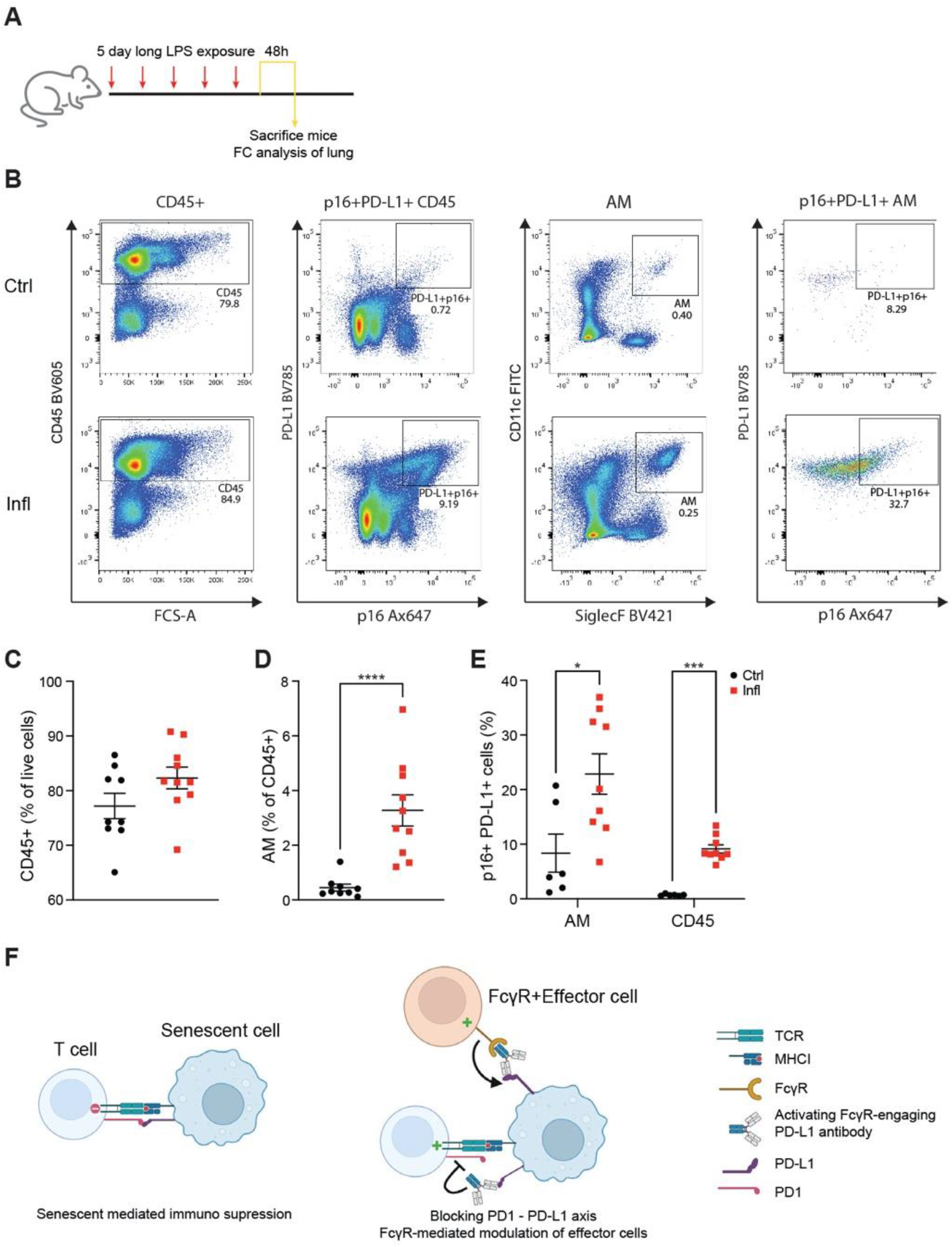
Anti-PD-L1 antibody treatment depletes p16, PD-L1 positive cells *in vivo*. **(A)** Experimental setup: mice were exposed daily to either PBS (Ctrl) or LPS (Infl) inhalations for 5 days before analysis of mice lungs 48h after the last inhalation. **(B)** Representative flow cytometry plots showing the gating strategy for p16^+^ PD-L1^+^ cells from immune cells (CD45^+^) and Alveolar Macrophage (AM) population. **(C-E)** Flow cytometry analysis of (C) CD45^+^, (D) AM, and (E) p16^+^PD-L1^+^ cells within CD45^+^ and AM populations in mice treated as in A. **(F)** Scheme of anti-PD-L1 antibody treatment and its effect on senescent cell immune surveillance.

